# MRIQC: Advancing the Automatic Prediction of Image Quality in MRI from Unseen Sites

**DOI:** 10.1101/111294

**Authors:** Oscar Esteban, Daniel Birman, Marie Schaer, Oluwasanmi O. Koyejo, Russell A. Poldrack, Krzysztof J. Gorgolewski

## Abstract

Quality control of MRI is essential for excluding problematic acquisitions and avoiding bias in subsequent image processing and analysis. Visual inspection is subjective and impractical for large scale datasets. Although automated quality assessments have been demonstrated on single-site datasets, it is unclear that solutions can generalize to unseen data acquired at new sites. Here, we introduce the MRI Quality Control tool (*MRIQC*), a tool for extracting quality measures and fitting a binary (accept/exclude) classifier. Our tool can be run both locally and as a free online service via the OpenNeuro.org portal. The classifier is trained on a publicly available, multi-site dataset (17 sites, N=1102). We perform model selection evaluating different normalization and feature exclusion approaches aimed at maximizing across-site generalization and estimate an accuracy of 76%*±*13% on new sites, using leave-one-site-out cross-validation. We confirm that result on a held-out dataset (2 sites, N=265) also obtaining a 76% accuracy. Even though the performance of the trained classifier is statistically above chance, we show that it is susceptible to site effects and unable to account for artifacts specific to new sites. MRIQC performs with high accuracy in intra-site prediction, but performance on unseen sites leaves space for improvement which might require more labeled data and new approaches to the between-site variability. Overcoming these limitations is crucial for a more objective quality assessment of neuroimaging data, and to enable the analysis of extremely large and multi-site samples.

## Introduction

Image analysis can lead to erroneous conclusions when the original data are of low quality (e.g. [1–4]). MRI images are unlikely to be artifact-free, and assessing their quality has long been a challenging issue [5]. Traditionally, all images in a sample under analysis are visually inspected by one or more experts, and those showing an insufficient level of quality are excluded (some examples are given in Fig 1). Visual assessment is time consuming and prone to variability due to inter-rater differences (see Fig 2), as well as intra-rater differences arising from factors such as practice or fatigue. An additional concern is that some artifacts evade human detection entirely [7], such as those due to improper choice of acquisition parameters. Even though magnetic resonance (MR) systems undergo periodic inspections and service, some machine-related artifacts persist unnoticed due to lenient vendor quality checks and drift from the system calibration settings. In our experience, automated quality control (QC) protocols help detect these issues early in the processing stream. Furthermore, the current trend towards acquiring very large samples across multiple scanning sites [8–10] introduces additional concerns. These large scale imaging efforts render the visual inspection of every image infeasible and add the possibility of between-site variability. Therefore, there is a need for fully-automated, robust and minimally biased QC protocols. These properties are difficult to achieve for three reasons: 1) the absence of a “gold standard” impedes the definition of relevant quality metrics; 2) human experts introduce biases with their visual assessment; and 3) cross-study and inter-site acquisition differences introduce uncharacterized variability. Machine-specific artifacts have generally been tracked down in a quantitative manner using phantoms [11]. However, many forms of image

**Figure 1.**
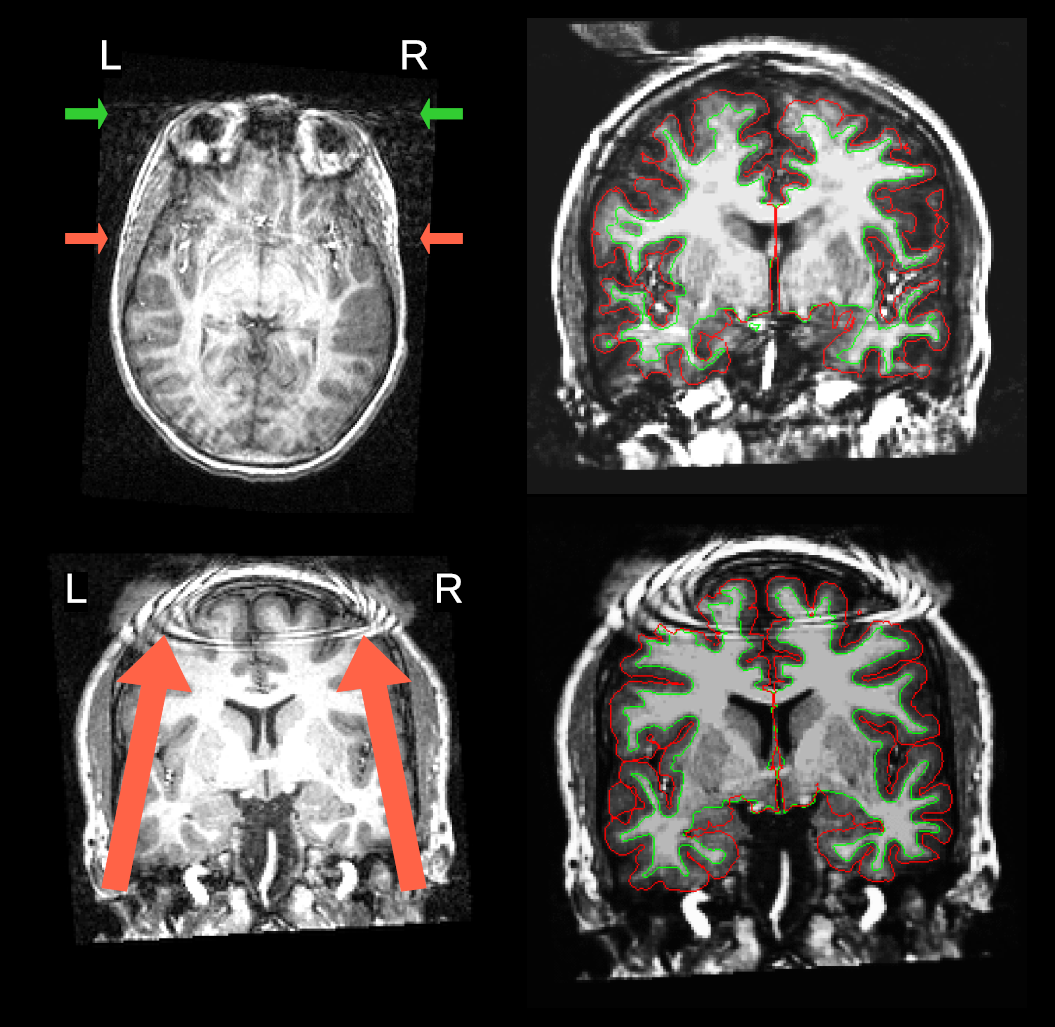
Visual assessment of MR scans. Two images with prominent artifacts from the Autism Brain Imaging Data Exchange (ABIDE) dataset are presented on the left. An example scan (top) is shown with severe motion artifacts. The arrows point to signal spillover through the phase-encoding axis (right-to-left –RL–) due to eye movements (green) and vessel pulsations (red). A second example scan (bottom) shows severe coil artifacts. On the right, the panel displays one representative image frame extracted from the animations corresponding to the subjects presented on the left, as they are inspected by the raters during the animation. This figure caption is extended in Figure SI1.

**Figure 2.**
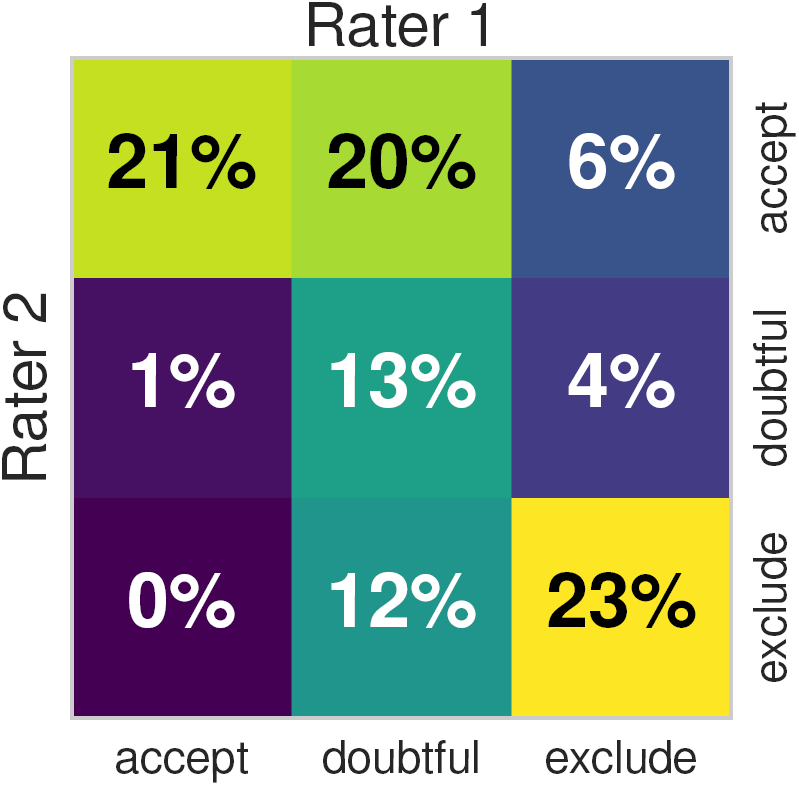
Inter-rater variability. The heatmap on the left shows the overlap of the quality labels assigned by two different domain experts on 100 data points of the ABIDE dataset, using the protocol described in section Labeling protocol. We also compute the Cohen’s Kappa index of both ratings, and obtain a value of *κ*=0.39. Using the table for interpretation of *κ* by Viera et al. [6], the agreement of both raters is “fair” to “moderate”. When the labels are binarized by mapping “doubtful” and “accept” to a single “good” label, the agreement increases to *κ*=0.51 (“moderate”). The “fair” to “moderate” agreement of observers demonstrates a substantial inter-rater variability. The inter- and intra-rater variabilities translate into the problem as *class-noise* since a fair amount of data points are assigned noisy labels that are not consistent with the labels assigned on the rest of the dataset. An extended investigation of the inter- and intra-rater variabilities is presented in SI: Impact of the labeling protocol and variability sources. degradation are participant-specific (e.g. the examples in Fig 1) or arise from practical settings (for instance, aliasing produced by the use of headsets during acquisition).

The automated quality control of magnetic resonance imaging (MRI) has long been an open issue. Woodard and Carley-Spencer [12] conducted one of the earliest evaluations on a large dataset of 1001 T1-weighted (T1w) MR images from 143 participants. They defined a set of 239 *no-reference* (i.e. no ground-truth of the same image without degradation exists) image-quality metrics (IQMs). The IQMs belonged to two families depending on whether they were derived from Natural Scene Statistics or quality indices defined by the JPEG consortium. The IQMs were calculated on image pairs with and without several synthetic distortions. In an analysis of variance, some IQMs from both families reliably discriminated among undistorted images, noisy images, and images distorted by intensity non-uniformity (INU). Mortamet et al. [13] proposed two quality indices focused on detecting artifacts in the air region surrounding the head, and analyzing the goodness-of-fit of a model for the background noise. One principle underlying their proposal is that most of the artifact signal propagates over the image and into the background. They applied these two IQMs on 749 T1w scans from the Alzheimer’s Disease Neuroimaging Initiative (ADNI) dataset. By defining cutoff thresholds for the two IQMs, they assigned the images high or low quality labels, and compared this classification to a manual assessment. They concluded that more specific research was required to determine these thresholds and generalize them to different datasets. However, many potential sources of uncontrolled variability exist between studies and sites, including MRI protocols (scanner manufacturer, MR sequence parameters, etc.), scanning settings, participant instructions, inclusion criteria, etc. For these reasons, the thresholds they proposed on their IQMs are unlikely to generalize beyond the ADNI dataset.

Later efforts to develop IQMs appropriate for MRI include the Quality Assessment Protocol (QAP), and the UK Biobank [14]. MRIQC extends the list of IQMs from the QAP, which was constructed from a careful review of the MRI and medical imaging literature [15]. Recently, Pizarro et al. [16] proposed the use of a support-vector machine classifier (SVC) trained on 1457 structural MRI images acquired in one site

The hypothesis motivating the present study is that the quality ratings of an expert on previously unseen datasets (with dataset-specific scanning parameters) can be predicted with a supervised learning approach that uses a number of IQMs as features. The first limitation we shall encounter when trying to answer this question is the inter-site variability of features extracted from MRI. Many efforts have been devoted to the normalization across sites of the intensities of T1w MRI [17]. Particularly, this inter-site variability renders as a *batch effect* problem in our derived IQMs (Fig 3), a concept arising from the analysis of gene-expression arrays [18]. Furthermore, the inherent subjectivity of the ratings done by experts, the difficulty of minimizing inter-rater variability and the particular labeling protocol utilized all introduce *class-noise* in the labels manually assigned. To demonstrate that the trained classifier correctly predicts the quality of new data, we used two unrelated datasets to configure the training and a held-out (test) datasets [19]. We first select the best performing model on the training dataset using a grid strategy in a nested cross-validation setup. We use the ABIDE dataset [9] as a training set because data are acquired in 17 different scanning sites with varying acquisition parameters (Table 1). These data show great variability in terms of imaging settings and parameters, which accurately represents the heterogeneity of real data and introduces the *batch effect* into modeling. The best performing classifier is then trained on the full ABIDE dataset and tested on the held-out dataset (*DS030* [20]), which is completely independent of ABIDE, to evaluate prediction on new sites.

**Figure 3.**
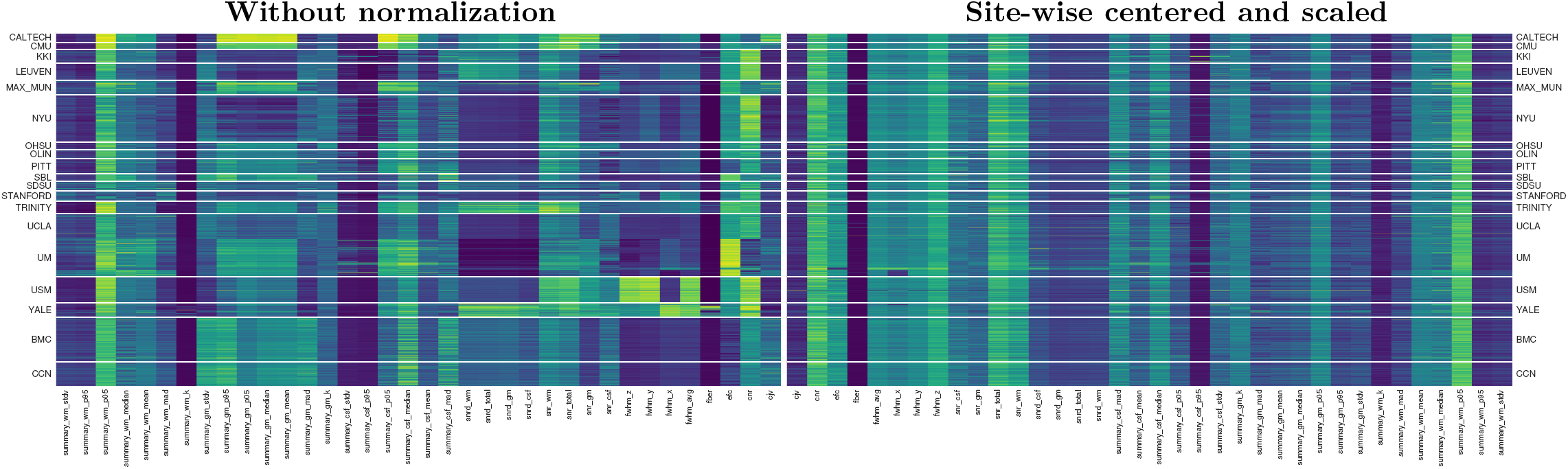
Inter-site variability renders as a *batch effect* on the calculated IQMs. These plots display features extracted by MRIQC (columns) of all participants (rows), clustered by site (17 centers from the ABIDE datasets, plus the two centers where DS030 was acquired –”BMC” and “CCN”–). The plot of original features (left panel) shows how they can easily be clustered by the site they belong to. After site-wise normalization including centering and scaling within site (right), the measures are more homogeneous across sites. Features are represented in arbitrary units. For better interpretation, the features-axis (*x*) has been mirrored between plots. with constant scanning parameters. They proposed three volumetric features and three features targeting particular artifacts. The volumetric features were the normalized histogram, the tissue-wise histogram and the ratio of the modes of gray matter (GM) and white matter (WM). The artifacts addressed were the eye motion spillover in the anterior-to-posterior phase-encoding direction, the head-motion spillover over the nasio-cerebellum axis (which they call *ringing artifact*) and the so-called wrap-around (which they refer to as *aliasing artifact*). They reported a prediction accuracy around 80%, assessed using 10-fold cross-validation. These previous efforts succeeded in showing that automating quality ratings of T1w MRI scans is possible. However, they did not achieve generalization across multi-site datasets.

**Table 1.**
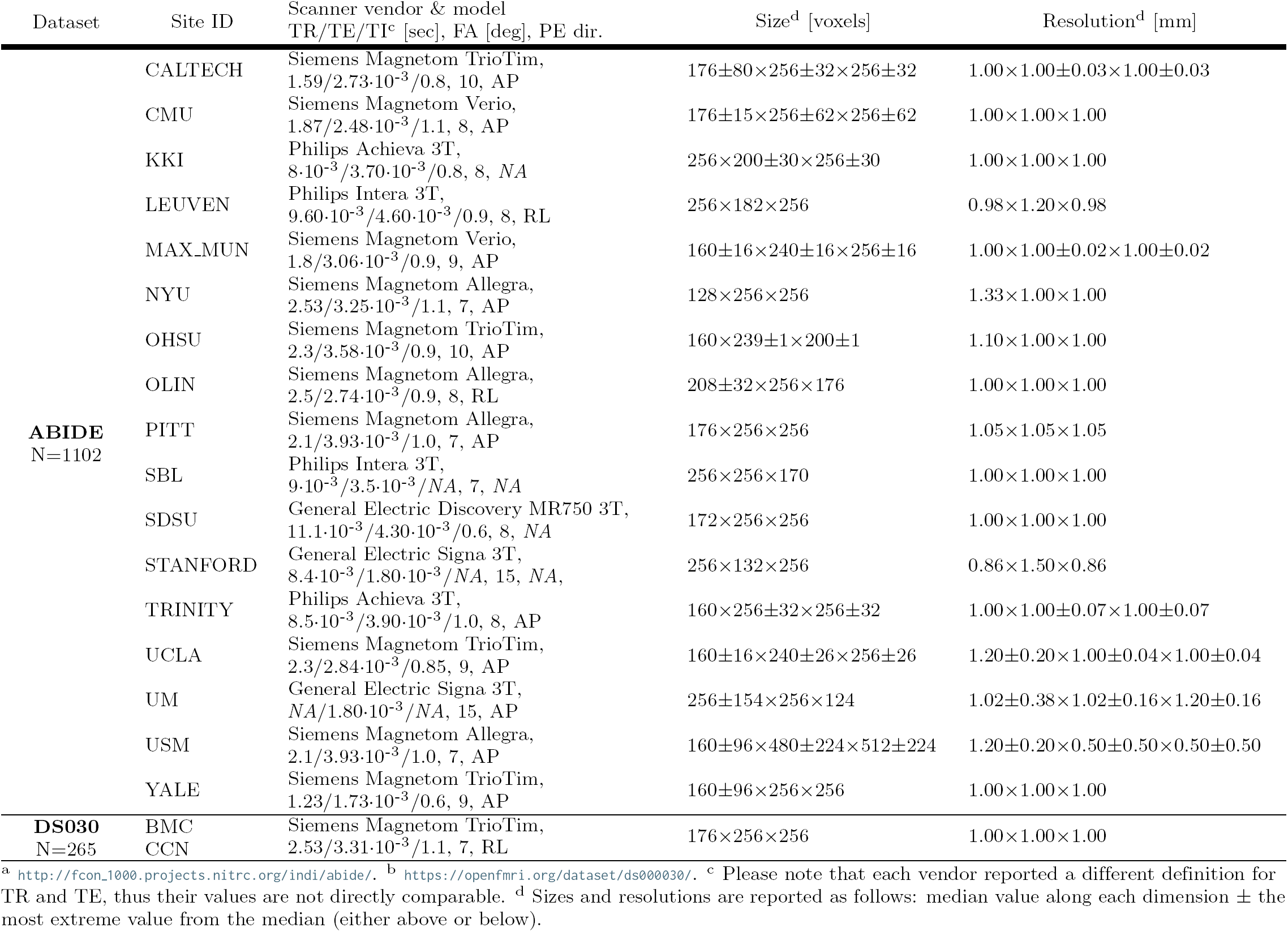
Summary table of the train and test datasets. The ABIDE dataset is publicly available^a^, and contains images acquired at 17 sites, with a diverse set of acquisition settings and parameters. This heterogeneity makes it a good candidate to train machine learning models that can generalize well to novel samples from new sites. We selected *DS030* [20] from OpenfMRI^b^ as held-out dataset to evaluate the performance on data unrelated to the training set. A table summarizing the heterogeneity of parameters within the ABIDE dataset and also *DS030* is provided with the supplemental materials (Table SI1).

| Dataset | Site ID | Scanner vendor & model<br>TR/TE/TI<sup>c</sup> [sec], FA [deg], PE dir. | Size<sup>d</sup> [voxels] | Resolution<sup>d</sup> [mm] |
| --- | --- | --- | --- | --- |
| ABIDE<br>N=1102 | CALTECH | Siemens Magnetom TrioTim,<br>1.59/2.73·10<sup>-3</sup>/0.8, 10, AP | 176±80×256±32×256±32 | 1.00×1.00±0.03×1.00±0.03 |
|  | CMU | Siemens Magnetom Verio,<br>1.87/2.48·10<sup>-3</sup>/1.1, 8, AP | 176±15×256±62×256±62 | 1.00×1.00×1.00 |
|  | KKI | Philips Achieva 3T,<br>8·10<sup>-3</sup>/3.70·10<sup>-3</sup>/0.8, 8, NA | 256×200±30×256±30 | 1.00×1.00×1.00 |
|  | LEUVEN | Philips Intera 3T,<br>9.60·10<sup>-3</sup>/4.60·10<sup>-3</sup>/0.9, 8, RL | 256×182×256 | 0.98×1.20×0.98 |
|  | MAX_MUN | Siemens Magnetom Verio,<br>1.8/3.06·10<sup>-3</sup>/0.9, 9, AP | 160±16×240±16×256±16 | 1.00×1.00±0.02×1.00±0.02 |
|  | NYU | Siemens Magnetom Allegra,<br>2.53/3.25·10<sup>-3</sup>/1.1, 7, AP | 128×256×256 | 1.33×1.00×1.00 |
|  | OHSU | Siemens Magnetom TrioTim,<br>2.3/3.58·10<sup>-3</sup>/0.9, 10, AP | 160×239±1×200±1 | 1.10×1.00×1.00 |
|  | OLIN | Siemens Magnetom Allegra,<br>2.5/2.74·10<sup>-3</sup>/0.9, 8, RL | 208±32×256×176 | 1.00×1.00×1.00 |
|  | PITT | Siemens Magnetom Allegra,<br>2.1/3.93·10<sup>-3</sup>/1.0, 7, AP | 176×256×256 | 1.05×1.05×1.05 |
|  | SBL | Philips Intera 3T,<br>9·10<sup>-3</sup>/3.5·10<sup>-3</sup>/NA, 7, NA | 256×256×170 | 1.00×1.00×1.00 |
|  | SDSU | General Electric Discovery MR750 3T,<br>11.1·10<sup>-3</sup>/4.30·10<sup>-3</sup>/0.6, 8, NA | 172×256×256 | 1.00×1.00×1.00 |
|  | STANFORD | General Electric Signa 3T,<br>8.4·10<sup>-3</sup>/1.80·10<sup>-3</sup>/NA, 15, NA, | 256×132×256 | 0.86×1.50×0.86 |
|  | TRINITY | Philips Achieva 3T,<br>8.5·10<sup>-3</sup>/3.90·10<sup>-3</sup>/1.0, 8, AP | 160×256±32×256±32 | 1.00×1.00±0.07×1.00±0.07 |
|  | UCLA | Siemens Magnetom TrioTim,<br>2.3/2.84·10<sup>-3</sup>/0.85, 9, AP | 160±16×240±26×256±26 | 1.20±0.20×1.00±0.04×1.00±0.04 |
|  | UM | General Electric Signa 3T,<br>NA/1.80·10<sup>-3</sup>/NA, 15, AP | 256±154×256×124 | 1.02±0.38×1.02±0.16×1.20±0.16 |
|  | USM | Siemens Magnetom Allegra,<br>2.1/3.93·10<sup>-3</sup>/1.0, 7, AP | 160±96×480±224×512±224 | 1.20±0.20×0.50±0.50×0.50±0.50 |
|  | YALE | Siemens Magnetom TrioTim,<br>1.23/1.73·10<sup>-3</sup>/0.6, 9, AP | 160±96×256×256 | 1.00×1.00×1.00 |
| DS030<br>N=265 | BMC | Siemens Magnetom TrioTim,<br>2.53/3.31·10<sup>-3</sup>/1.1, 7, RL | 176×256×256 | 1.00×1.00×1.00 |
|  | CCN |  |  |  |

<sup>a</sup> http://fcon.1000.projects.nitrc.org/indi/abide/. <sup>b</sup> https://openfmri.org/dataset/ds000030/. <sup>c</sup> Please note that each vendor reported a different definition for TR and TE, thus their values are not directly comparable. <sup>d</sup> Sizes and resolutions are reported as follows: median value along each dimension ± the most extreme value from the median (either above or below).

The contributions of this work are summarized as follows. First, we release an MRI quality control tool called MRIQC (described in The MRIQC tool) to extract a vector of 64 IQMs (Table 2) per input image (Extracting the Image Quality Metrics). Second, MRIQC includes a visual reporting system (described in the Reports for visual assessment section) to ease the manual investigation of potential quality issues. These visual reports allow researchers to quickly evaluate the cases flagged by the MRIQC classifier or visually identify potential images to be excluded by looking at the group distributions of IQMs. Finally, we describe the results of the pre-registered report corresponding to this study (see Software and data availability) on the feasibility of automatic quality rating and the implications of the inter-site variability of IQMs (sections Supervised classification and Results).

## Materials and Methods

### Training and test datasets

A total of 1367 T1w scans are used as training (1102 from ABIDE) and test (265 from *DS030*) samples. These datasets are intentionally selected for their heterogeneity to match the purpose of the study. A brief summary illustrating the diversity of acquisition parameters is presented in Table 1, and a full-detail table in Table SI1.

#### Labeling protocol

Based on our experience and minimizing the time-cost of inspecting each of the 1367 images, we designed an agile labeling protocol as follows. The experts visualize an animated GIF (graphics interchange format) video sequentially showing coronal slices (in anterior to posterior ordering) of the image under assessment. Each animation has a duration of around 20s (see Software and data availability). During the visualization, the rater assesses the overall quality of the image. Raters were asked to assign a quality label (“exclude”, “doubtful” or “accept”) based on their experience after inspection of each animation. The animation is replayed in loop until the rater makes a decision.

The labeling process is aided by surface reconstructions, using the so-called *white* (WM-GM interface) and the *pial* (delineating the outer interface of the cortex) surfaces as visual cues for the rater. The *white* and *pial* contours are used as evaluation surrogates, given that “exclude” images usually exhibit imperfections and inaccuracies on these surfaces. When the expert finds general quality issues or the reconstructed surfaces reveal more specific artifacts, the “exclude” label is assigned and the rater notes a brief description, for example: “low signal-to-nose ratio (SNR)”, “poor image contrast”, “ringing artifacts”, “head motion”, etc. We utilize *FreeSurfer 5.3.0* [21] to reconstruct the surfaces. *FreeSurfer* has been recently reported as a good quality proxy to assess T1w images [22]. For run-time considerations, and to avoid circular evaluations of *FreeSurfer*, this tool is not used in the MRIQC workflow (see The MRIQC tool section).

The first rater (MS) assessed 601 images of ABIDE, covering *∼*55% of the dataset. The second rater (DB) also assessed 601 images from the ABIDE dataset, and all the 265 images of the *DS030* dataset. Since both raters covered more than half of ABIDE, one hundred images of the dataset were rated by both experts. Such overlap of assessments enables the characterization of inter-rater variability (Fig 2) using those images with double ratings. For training, we randomly draw fifty ratings of each expert (without replacement) from the one hundred data points assessed twice. The images of the ABIDE dataset were randomized before splitting by rater. Additionally, the participant identifier was blinded in the animations. Participants in both datasets were randomized before rating to avoid inducing site-specific *class-noise*. Finally, the ABIDE rating process yielded a balance of 337/352/412 exclude/doubtful/accept data points (31%/32%/37%, see SI: Impact of the labeling protocol and variability sources). Balances for *DS030* are 75/145/45 (28%/55%/17%).

### Software instruments and calculation of the IQMs

#### The MRIQC tool

MRIQC is an open-source project, developed under the following software engineering principles. 1) *Modularity and integrability*: MRIQC implements a *nipype* [23] workflow (see Fig 4) to integrate modular sub-workflows that rely upon third party software toolboxes such as *FSL* [24], *ANTs* [25] and *AFNI* [26]. 2) *Minimal preprocessing*: the workflow should be as minimal as possible to estimate the IQMs. 3) *Interoperability and standards*: MRIQC is compatible with input data formatted according to the the Brain Imaging Data Structure (BIDS, [27]) standard, and the software itself follows the BIDS Apps [28] standard. For more information on how to convert data to BIDS and run MRIQC, see SI: Converting datasets into BIDS and SI: Running MRIQC respectively. 4) *Reliability and robustness*: the software undergoes frequent vetting sprints by testing its robustness against data variability (acquisition parameters, physiological differences, etc.) using images from the OpenfMRI resource. Reliability is checked and tracked with a continuous integration service.

**Figure 4.**
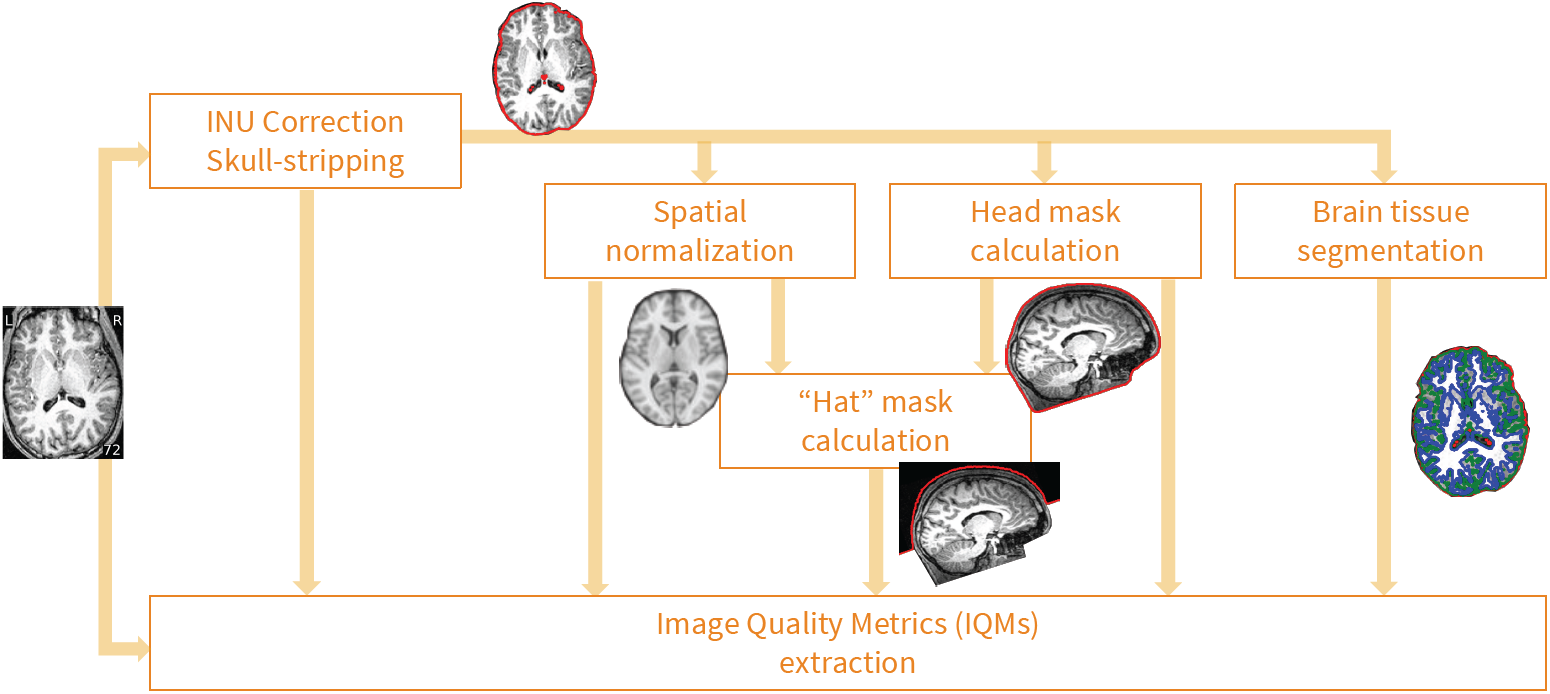
MRIQC’s processing data flow. Images undergo a minimal processing pipeline to obtain the necessary corrected images and masks required for the computation of the IQMs.

#### Extracting the Image Quality Metrics

The final steps of MRIQC’s workflow compute the different IQMs, and generate a summary JSON file per subject. The IQMs can be grouped in four broad categories (see Table 2), providing a vector of 64 features per anatomical image. Some measures characterize the impact of noise and/or evaluate the fitness of a noise model. A second family of measures uses information theory and prescribed masks to evaluate the spatial distribution of information. A third family of measures looks for the presence and impact of particular artifacts. Specifically, the INU artifact, and the signal leakage due to rapid motion (e.g. eyes motion or blood vessel pulsation) are identified. Finally, some measures that do not fit within the previous categories characterize the statistical properties of tissue distributions, volume overlap of tissues with respect to the volumes projected from MNI space, the sharpness/blurriness of the images, etc. The ABIDE and *DS030* datasets are processed utilizing *Singularity* [29] (see SI: Reproducing the experiments).

**Table 2.** Summary table of IQMs. The 14 IQMs spawn a vector of 64 features per anatomical image on which the classifier is learned and tested.

| Measures based on noise measurements |  |
| --- | --- |
| CJV | The coefficient of joint variation of GM and WM was proposed as objective function by Ganzetti et al. [30] for the optimization of INU correction algorithms. Higher values are related to the presence of heavy head motion and large INU artifacts. |
| CNR | The contrast-to-noise ratio [31] is an extension of the SNR calculation to evaluate how separated the tissue distributions of GM and WM are. Higher values indicate better quality. |
| SNR | MRIQC includes the signal-to-noise ratio calculation proposed by Dietrich et al. [32], using the air background as noise reference. Additionally, for images that have undergone some noise reduction processing, or the more complex noise realizations of current parallel acquisitions, a simplified calculation using the within tissue variance is also provided. |
| QI<sub>2</sub> | The second quality index of [13] is a calculation of the goodness-of-fit of a $\chi^2$ distribution on the air mask, once the artifactual intensities detected for computing the QI<sub>1</sub> index have been removed. The description of the QI<sub>1</sub> is found below. |
| Measures based on information theory |  |
| EFC | The entropy-focus criterion [33] uses the Shannon entropy of voxel intensities as an indication of ghosting and blurring induced by head motion. Lower values are better. |
| FBER | The foreground-background energy ratio [15] is calculated as the mean energy of image values within the head relative the mean energy of image values in the air mask. Consequently, higher values are better. |
| Measures targeting specific artifacts |  |
| INU | MRIQC measures the location and spread of the bias field extracted estimated by the intensity non-uniformity correction. The smaller spreads located around 1.0 are better. |
| QI<sub>1</sub> | The first quality index of [13] measures the amount of artifactual intensities in the air surrounding the head above the nasio-cerebellar axis. The smaller QI<sub>1</sub>, the better. |
| WM2MAX | The white-matter to maximum intensity ratio is the median intensity within the WM mask over the 95% percentile of the full intensity distribution, that captures the existence of long tails due to hyper-intensity of the carotid vessels and fat. Values should be around the interval [0.6, 0.8]. |
| Other measures |  |
| FWHM | The full-width half-maximum [34] is an estimation of the blurriness of the image using AFNI's 3dFWHMx. Smaller is better. |
| ICVs | Estimation of the intracranial volume of each tissue calculated on the FSL FAST's segmentation. Normative values fall around 20%, 45% and 35% for cerebrospinal fluid (CSF), WM and GM, respectively. |
| rPVE | The residual partial volume effect feature is a tissue-wise sum of partial volumes that fall in the range [5%-95%] of the total volume of a pixel, computed on the partial volume maps generated by FSL FAST. Smaller residual partial volume effects (rPVEs) are better. |
| SSTATs | Several summary statistics (mean, standard deviation, percentiles 5% and 95%, and kurtosis) are computed within the following regions of interest: background, CSF, WM, and GM. |
| TPMs | Overlap of tissue probability maps estimated from the image and the corresponding maps from the ICBM nonlinear-asymmetric 2009c template [35]. |

#### Reports for visual assessment

In order to ease the screening process of individual images, MRIQC generates individual reports with mosaic views of a number of cutting planes and supporting information (for example, segmentation contours). The most straightforward use-case is the visualization of those images flagged by the classifier. After the extraction of IQMs from all images in the sample, a group report is generated (Fig 5). The group report shows a scatter plot for each of the IQMs, so it is particularly easy to notice cases that are outliers for each metric. The plots are interactive, such that clicking on any particular sample opens up the corresponding individual report of that case. Examples of group and individual reports for the ABIDE dataset are available online at mriqc.org.

**Figure 5.**
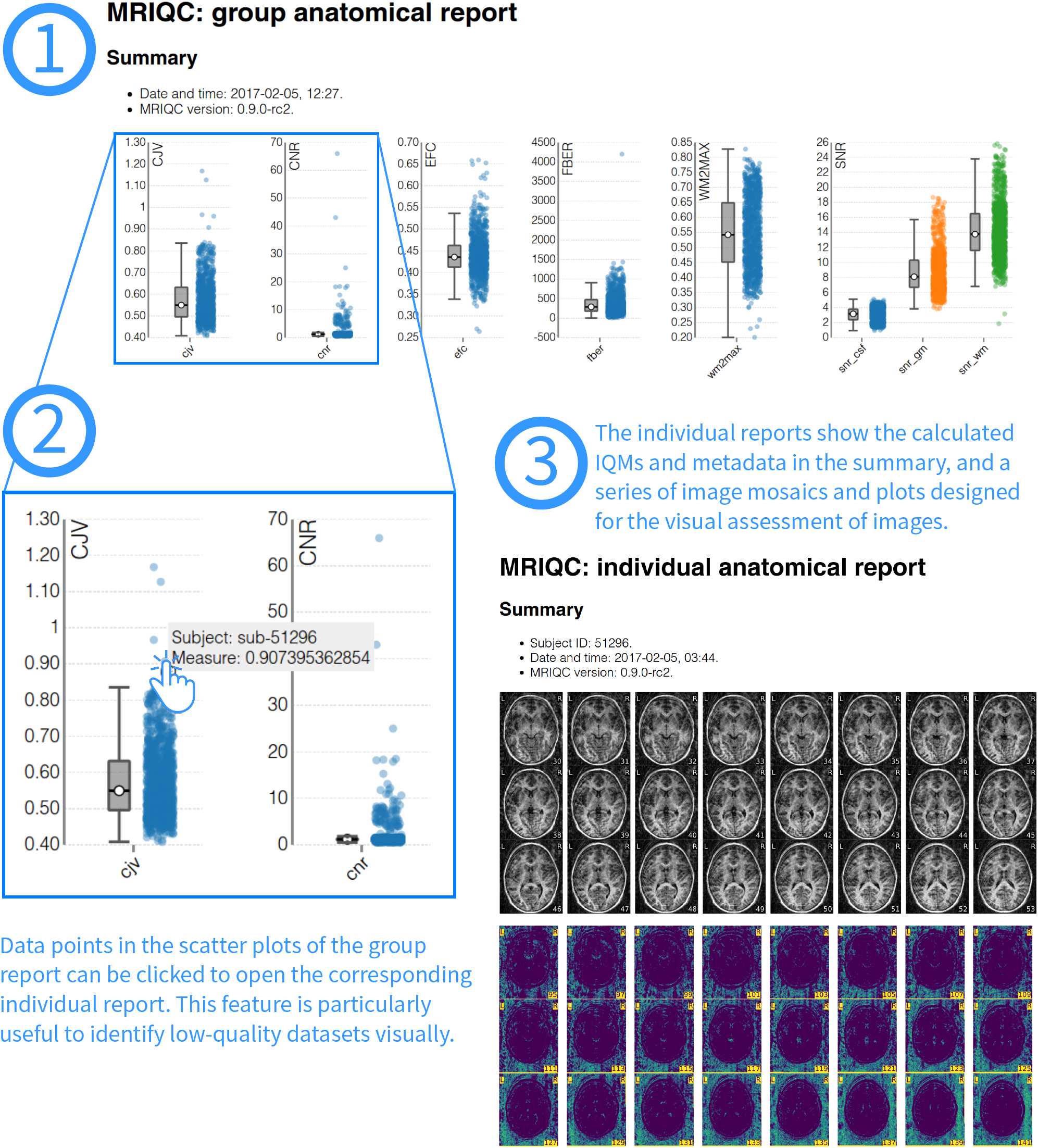
Visual reports. MRIQC generates one individual report per subject in the input folder and one group report including all subjects. To visually assess MRI samples, the first step (1) is opening the group report. This report shows boxplots and strip-plots for each of the IQMs. Looking at the distribution, it is possible to find images that potentially show low-quality as they are generally reflected as outliers in one or more strip-plots. For instance, in (2) hovering a suspicious sample within the coefficient of joint variation (CJV) plot, the subject identifier is presented (“sub-51296”). Clicking on that sample will open the individual report for that specific subject (3). This particular example of individual report is available online at https://web.stanford.edu/group/poldracklab/mriqc/reports/sub-51296T1w.html.

### Supervised classification

We propose a supervised classification framework using the ABIDE dataset as the training set and *DS030* as the held-out dataset. Both datasets are rated as described in Labeling protocol by two experts. Labels are binarized mapping the “doubtful” and “accept” labels to a single “accept” quality rating. We use cross-validation to evaluate the performance of the models. Prior to model selection using cross-validation, we first investigate the appropriate cross-validation design most adequate to pick on the *batch effects*, avoiding an overly optimistic evaluation of the performance (see What data split should be used in cross-validation?). We then present a two-step approach to predicting the quality labels of the held-out dataset. First, we perform a preliminary evaluation using nested cross-validation utilizing only the ABIDE dataset (see Step 1: Tested models and selection) to choose the best performing model. Then, we optimize it in a refined grid of hyper-parameters with a single-loop cross-validation on the ABIDE dataset. Finally, the model is evaluated using the held-out dataset (see Step 2: Validation on the held-out dataset). The cross-validation workflows are built upon *scikit-learn* [36] and run utilizing *Singularity* [29] (see SI: Reproducing the experiments).

#### Step 1: Tested models and selection

Based on the number of features (64) and training data available (*∼* 1100 data points), we compare two families of classifiers: SVCs and random forests classifiers (RFCs). We evaluate several preprocessing alternatives to overcome the *batch effects*. In order to deal with the class-imbalance, we also evaluate all models with and without class weighting during resampling. When enabled, weighting is inversely proportional to class frequencies in the input data.

#### The support-vector machine classifier (SVC)

A support-vector machine [37] finds a hyperplane in the high-dimensional space of the features that robustly separates the classes of interest. The SVC then uses the hyperplane to decide the class that is assigned to new samples in the space of features. Two hyper-parameters define the support-vector machine algorithm: a kernel function that defines the similarity between data points to ultimately compute a distance to the hyperplane, and a regularization weight *C*. In particular, we analyzed here the linear SVC implementation (as of now, “SVC-lin”) and the one based on radial basis functions (denoted by “SVC-rbf”). During model selection, we evaluated the regularization weight *C* of both SVCs and the *γ* parameter (kernel width) particular to the SVC-rbf.

#### The random forests classifier (RFC)

Random forests [38] are a nonparametric ensemble learning method that builds multiple decision trees. The RFC assigns to each new sample the mode of the predicted classes of all decision trees in the ensemble. In this case, random forests are driven by a larger number of hyper-parameters. Particularly, we analyze the number of decision trees, the maximum tree-depth, the minimum number of samples per split, and the minimum node size.

#### Objective function

The performance of each given model and parameter selection can be quantified with different metrics. Given the imbalance of positive and negative cases –with lower prevalence of “exclude” samples–, we select the area under the curve (AUC) of the receiver-operator characteristic as objective score. Additionally, we report the classification accuracy (ACC) as implemented in *scikit-learn* (see Equation SI1).

#### Preprocessing

In order to address the *batch effect* and build models more robust to this problem, we include three preprocessing steps in the framework. The first preprocessing step is a *site-wise* normalization of features. For robustness, this normalization calculates a center (as the median feature value) and a spread (as the interquartile range) per feature for demeaning and scaling data. This filter can center only, scale only or perform both centering and scaling.

The second preprocessing step available is a dimensionality reduction filter excluding features highly predictive of the site of origin of data points. To do so, we fit a classifier based on extremely randomized trees [39], where the variables are the features and the responses are the sites of acquisition. We iteratively fit the classifier and remove the feature most predictive of the site at each step, until certain convergence criteria is met (either a maximum number of features to remove is reached or the performance of the classifier is very low and thus the remaining features do not predict the site at all).

Finally, a third preprocessing step implements the Winnow algorithm [40] using extremely randomized trees in a similar way to the previous filter, but comparing features to a synthetic, randomly-generated feature. This feature selection filter removes those IQMs below a certain SNR level.

All the hyper-parameters (normalization centering and/or scaling and the two feature elimination algorithms) can be switched on and off during model selection. Finally, they are optimized in a cross-validation framework.

#### Cross-validation and nested cross-validation

Cross-validation is a model selection and validation technique that can be robust to data inhomogeneities [41] with the appropriate choice of the data split scheme. We use nested cross-validation, which divides the process into two validation loops: an inner loop for selecting the best model and hyper-parameters, and an outer loop for evaluation. In cross-validation, data are split into a number of folds, each containing a training and a test set. For each fold, the classifier is trained on the first set and evaluated on the latter. When cross-validation is nested, the training set is split again into folds within the inner loop, and training/evaluation are performed to optimize the model parameters. Only the best performing model of the inner loop is then cross-validated in the outer loop. Models and their hyper-parameters are evaluated within the inner loop, optimizing for the best average AUC score.

#### Data split scheme

To prevent the inflation of evaluation scores due to *batch effects*, we defined a *leave-one-site-out (LoSo)* partition strategy. The LoSo split leaves out a whole site as test set at each cross-validation fold. Therefore, no knowledge of the test site is leaked into the training set (the remaining *N* − 1 sites). In a preliminary experiment (What data split should be used in cross-validation?) we justify the use of LoSo over a more standard repeated and stratified 10-fold. If a *batch effect* exists it will result in over-fitting on the training data compared to the unseen test data. The performance measured in the outer cross-validation loop on the ABIDE dataset will be higher than that evaluating the classifier on the held-out dataset, unrelated to ABIDE.

#### Feature ranking

One tool to improve the interpretability of the RFC is the calculation of feature rankings [38] by means of variable importance or Gini importance. Since we use *scikit-learn*, the implementation is based on Gini importance, defined for a single tree as the total decrease in node impurity weighted by the probability of reaching that node. We finally report the median feature importance over all trees of the ensemble.

#### Step 2: Validation on the held-out dataset

In the second step, we use the model selected in step 1, and trained on the full ABIDE dataset to evaluate the performance on the held-out dataset (*DS030*).

## Results

All images included in the selected datasets are processed with MRIQC. After extraction of the IQMs from the ABIDE dataset, a total of 1101 images have both quality ratings and quality features (one image of ABIDE is skull-stripped, thus it is not valid for the extraction of measures with MRIQC and was excluded). In the case of *DS030*, all the 265 T1w images have the necessary quality ratings and features.

### What data split should be used in cross-validation?

Before fitting any particular model to the IQMs, we identify the cross-validation design most appropriate for the application. We confirm that the *batch effects* are overlooked when using a 10-Fold cross-validation like [16], producing a biased estimation of the performance. To support that intuition we run four nested cross-validation experiments, with varying split strategies for the inner and outer loops. First, the nested cross-validation is performed on the ABIDE dataset. We use a randomized search (evaluating 50 models) for the inner loop. Second, the cross-validated inner model is fitted onto the whole ABIDE dataset. The result in Table 3 shows that the LoSo splitting has a closer train-test accuracy than 10-Fold cross-validation.

**Table 3.** Selecting the appropriate split strategy for cross-validation. The cross-validated area under the curve (AUC) and accuracy (ACC) scores calculated on the ABIDE dataset (train set) are less biased when LoSo is used to create the outer folds, as compares to the evaluation scores obtained in *DS030* (held-out set).

| Outer split | Inner split | ABIDE (train) | | DS030 (held-out) | | Bias ($\Delta$) | |
| --- | --- | --- | --- | --- | --- | --- | --- |
|  |  | AUC | ACC (%) | AUC | ACC (%) | AUC | ACC |
| 10-Fold | 5-Fold | .87±.04 | 83.75±3.6 | .68 | 75.7 | .19 | 7.0 |
| | LoSo | .86±.04 | 81.93±3.5 | .71 | 77.0 | .15 | 5.0 |
| LoSo | 5-Fold | .71±.15 | 75.96±16.8 | .68 | 76.2 | **.03** | **-0.2** |
| | LoSo | .71±.15 | 75.21±17.8 | .68 | 76.6 | **.03** | **-1.4** |

### Model evaluation and selection

Once the LoSo cross-validation scheme is selected, we use nested cross-validation to compare the three models investigated (*SVC-lin*, *SVC-rbf*, and *RFC*). Note that only the ABIDE dataset is used, therefore the *DS030* dataset is kept unseen during model selection. Again, the search strategy implemented for the inner cross-validation loop is a randomized search of 50 models. The best performing model is the RFC with all the optional preprocessing steps enabled. Therefore, the model includes the robust site-wise normalizer (with centering and scaling), the feature elimination based on predicting the site of origin and the Winnow-based feature selection. Fig 6 shows the AUC and ACC scores obtained for the three models evaluated, for each data split in the outer cross-validation loop.

**Figure 6.**
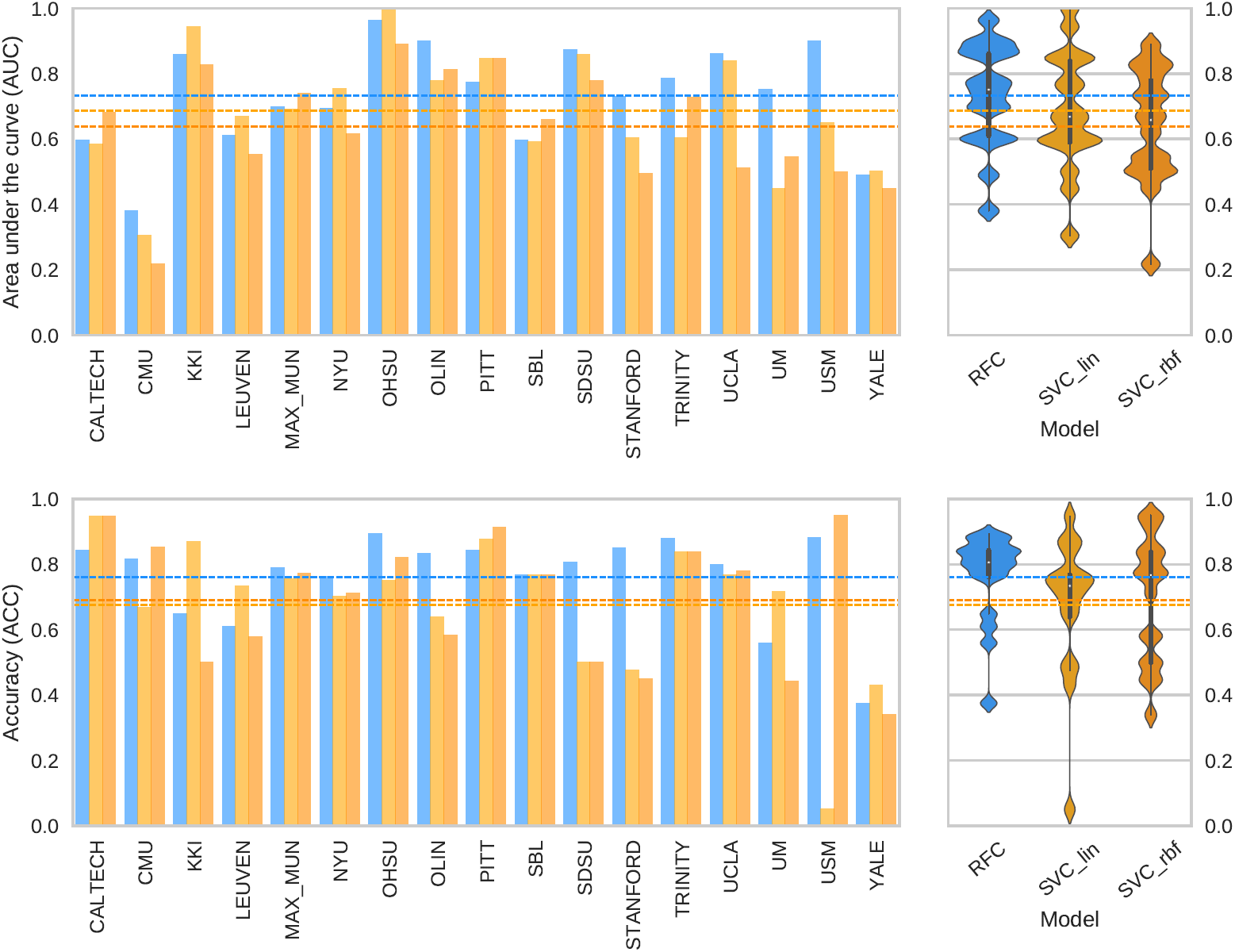
Nested cross-validation for model selection. The plots on the left represent the scores (AUC on top, ACC below) obtained in the outer loop of nested cross-validation, using the LoSo split. The plots show how certain sites are harder to predict than others. On the right, the corresponding violin plots that summarize the overall performance. In both plots, the dashed lines represent the averaged cross-validated performance for the three models: RFC (blue line, AUC=0.73*±*0.15, ACC=76.15%*±*13.38%), SVC-lin (light orange, AUC=0.68*±*0.18, ACC=67.54%*±*20.82%), and SVC-rbf (dark orange, AUC=0.64*±*0.17, ACC=69.05%*±*18.90%).

### Evaluation on held-out data

Finally, we run a non-nested cross-validation to find the best model and test it on the held-out dataset. In this case, we use a grid search strategy to evaluate all possible combinations of hyper-parameters (a total of 512 models). The specific grid we evaluate is available in the GitHub repository (mriqc/data/classifier settings.yml). In order to assess the above-chance accuracy performance, we run a permutation test [42] shuffling labels of both training and test sets at each repetition (1000 permutations). The evaluation on *DS030* is summarized on Table 4, and shows an AUC of 0.707, and ACC=76% (*p*=0.001). The performance is slightly higher than that (AUC/ACC=0.5/72%) of a naive classifier that labels all data points “accept”. The model selected includes the robust normalization (with both centering and scaling) and the site-prediction feature selection. The features finally selected are presented in the plot of feature importances of Fig 7 (panel A). The QI_2_ [13] is the most important feature, followed by background and WM tissue statistics. The recall (Equation SI2) is particularly low (0.28, Table 4B) and indicates over-fitting to the training set. To understand the problem, we visualized the images in the test set that were predicted “accept” but rated “exclude” by the expert, and found a signal ghost artifact in *∼*18% of the images in the test set that was not present in any image of the training set. Most of the images containing this artifact are rated as “exclude” by the expert, when the ghost artifact overlapped the cortical sheet of the temporal lobes. Some examples are reproduced in Fig 7B. To assess the performance in the absence of the ghosting artifact on *DS030*, we run the classifier trained on ABIDE on the test set after removing the images showing this artifact. The results of this exploratory analysis are presented in SI: The idiosyncratic ghost of DS030. Without the ghosting artifact, the performance improves to AUC/ACC=0.83/87%. The sensitivity to “exclude” images increases to 0.46. To ensure this increase of sensitivity is a direct consequence of the removal of the ghosting artifact, we re-run the nested cross-validation of the RFC model (see Model evaluation and selection) with a modification to report the recall. We obtain a value of 0.48 (*±*0.3), consistent with the previous result on the ghosting-free subsample of the *DS030* dataset.

**Figure 7.**
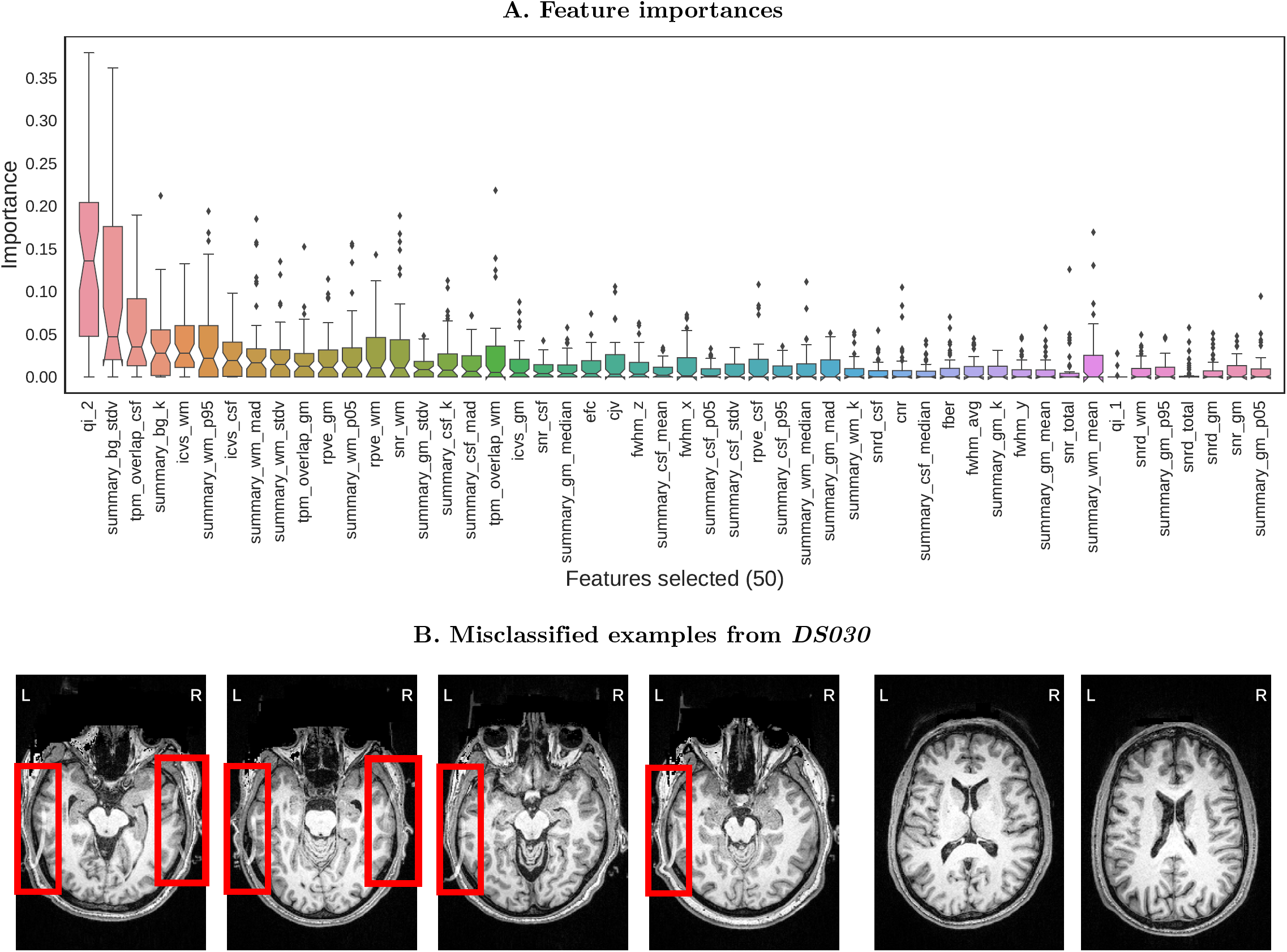
Evaluation on the held-out dataset. **A**. A total of 50 features are selected by the preprocessing steps. The features are ordered from highest median importance (the QI_2_ [13]) to lowest (percentile 5% of the intensities within the GM mask). The boxplots represent the distribution of importances of a given feature within all trees in the ensemble. **B**. (Left) Four different examples of false negatives of the *DS030* dataset. The red boxes indicate a ghosting artifact, present in more than 20% of the images. Only extreme cases where the ghost overlaps the cortical GM layer of the occipital lobes are presented. (Right) Two examples of false positives. The two examples are borderline cases that were rated as “doubtful”. Due to the intra- and inter-rater variabilities, some data points with poorer overall quality are rated just “doubtful”. These images demonstrate the effects of the noise in the quality labels.

**Table 4.** Evaluation on the held-out dataset. The model cross-validated on the ABIDE dataset performs with AUC=0.707 and ACC=76% on *DS030*. The confusion matrix shows an insensitivity of the classifier to many “exclude” cases (see the recall column for the “exclude” label in Table B).

| A. Confusion matrix |  |  |  |  | B. Performance report |  |  |  |  |
| --- | --- | --- | --- | --- | --- | --- | --- | --- | --- |
|  |  | Predicted |  |  |  |  |  |  |  |
|  |  | accept | reject | total |  | precision | recall | F1 score | support |
| True | accept | 180 | 10 | 190 | accept | 0.77 | 0.95 | 0.85 | 190 |
|  | reject | 54 | 21 | 75 | exclude | 0.68 | 0.28 | 0.40 | 75 |
|  | total | 234 | 31 |  | avg / total | 0.74 | 0.76 | 0.72 | 265 |

## Discussion

Quality control (QC) protocols identify faulty datasets that can bias analyses. We propose a quantitative approach to the QC of T1w MRI acquisitions of the brain. Human brain images can be degraded by various sources of artifacts related to the scanning device, session parameters, or the participants, themselves. Automating the QC process is particularly necessary for large scale, multi-site studies such as the UK Biobank. Previous efforts [12, 13, 16, 43] in the quantification of image quality are also based on no-reference image-quality metrics (IQMs), but did not attempt to solve the generalization of prediction to unseen samples from new sites.

In this work, we investigate the prediction of binary quality labels from a set of IQMs. As planned in the corresponding pre-registered report (see Software and data availability), we focus specifically on the generalization of prediction to image sets acquired in sites unseen by the classifier. Most of the IQMs used in this work and in previous literature [12, 13, 16, 43] are highly dependent on the specific acquisition parameters and particular scanning settings of each site. This inter-site variability is transfered into the IQMs producing *batch effects* [18] that impede the generalization of predictions to new sites (or “batches”). For these reasons, we pre-registered an experimental design based on a supervised learning framework using the ABIDE dataset as training set for its heterogeneity (acquired in 17 different sites), and one OpenfMRI dataset (*DS030*) as held-out dataset. We slightly deviate from the pre-registered design in minor details, for instance we do not use Bayesian estimation of hyper-parameters [44] since the use of grid search and randomized search are sufficient for the problem. Further deviations from the pre-registration are the increment on the number of IQMs used (we proposed 34, and use 64 here) and the final implementation of the MRIQC workflow. We also diverged in the finally applied Labeling protocol, since two different experts (instead of only one) manually rate a total of 1367 T1w images, and they did not revisit the “exclude” and “doubtful” cases using freeview. One expert evaluated 601 images belonging to the ABIDE dataset, and the second expert rated 601 images of ABIDE plus the full *DS030* (265 images). Thus, one hundred images of ABIDE selected randomly are rated by both experts. We utilize these overlapping ratings to investigate the inter-observer reliability using the Cohen’s Kappa (*κ*=0.39, “fair” agreement [6]).

When the ratings are binarized, *κ* increases to 0.51 (“moderate” [6]). This “fair” to “moderate” agreement unveils a second source of variance alongside the *batch effects*: the *class-noise* (or the variability in the assigned labels). We use cross-validation and nested cross-validation for model selection and evaluation. Before addressing the question of quality prediction, we first investigate the appropriate design of data splits for datasets showing *batch effects*. In section What data split should be used in cross-validation? we show that leave-one-site-out (LoSo) is a less optimistic and less biased design than a standard 10-fold split as in Pizarro et al. [16]. Once the cross-validation scheme is defined, we select the random forests classifier (RFC) model over two variants of the support-vector machine classifier (SVC) in a nested cross-validation scheme using only the ABIDE dataset. Finally, we select the final model and hyper-parameters in a non-nested cross-validation, train the model on the ABIDE dataset, and evaluate its performance on the held-out dataset (*DS030*). We obtain an area under the curve (AUC) score of *∼* 0.71 and an accuracy (ACC) score of *∼* 76%. We ensure the classifier is capturing the structure of quality labels from the data running a permutation test (*p*=0.001, 1000 permutations). The ultimately selected model includes the normalization of features (with both centering and scaling), and the feature elimination based on the site prediction (which removed 14 features highly correlated with the site of origin).

Intrigued by the poor sensitivity to positive (“exclude”) data points, we discover that *DS030* shows a systematic ghosting artifact in a substantial number of the images that is not present in any of the training examples (Fig 7B). Most of the images showing that artifact (except for a few where the ghost was present but did not overlap the cortical layer) are rated as “exclude” by the expert. In a subsequent exploratory analysis where we remove the data points presenting the artifact, we find that sensitivity to “exclude” cases rises from 0.28 to 0.46, and AUC/ACC improve from 0.71/76% to 0.83/87%. Therefore, the performance improves, albeit moderately. The sensitivity to “exclude” data points on the ghost-free test set is consistent to that estimated by means of nested cross-validation. On one hand, we argue that *DS030* is not a representative held-out dataset due to this structured artifact. On the other hand, it is likely that many scanning sites show idiosyncratic artifacts that are not present in our training set (ABIDE). We discuss this limitation with some others below.

We propose LoSo as cross-validation scheme in datasets showing *batch effects*. In the particular problem at hand, RFC outperforms SVCs and requires site-wise normalization of features to handle the heterogeneity of sites. The model selected in cross-validation also includes the feature selection by removing those features that best predicted the site of origin of samples. We interpret these results as a modest confirmation of the initial hypothesis, since the classifier captures the quality structure of features and it predicts the quality of the held-out dataset with above-chance accuracy. The performance we report is relatively low (even though not too far from recent studies using single-site data [16]), and we can hypothesize that it is easy to find a new sample that confuses the classifier just looking for particular artifacts not present in ABIDE and *DS030*. Therefore, the intent of generalization to new sites made in our initial hypothesis is only weakly confirmed at best.

One clear limitation of the presented classifier is the need for additional labeled data, acquired in new sites with scanners (vendor, model, software versions) and scanning parameters under-represented in the ABIDE dataset. Moreover, the images distributed under the ABIDE dataset have undergone a realignment process through resampling, that slightly modified the original intensity distributions and smoothed the images. One additional route to enhance the predictive power of the classifier is reducing the class-noise by refining the ratings done by the experts. Along the same lines, the ABIDE dataset could be augmented with images from new sites for which we correct MRIQC’s predictions *a posteriori*, and including these fixed data points within the training set. A similar approach to adding new sites to the training set would use techniques like label propagation [45], where only a random subset of the sample is manually rated and the labels are propagated to the remaining samples through an unsupervised clustering procedure. One more alternative to boost the prediction performance leverages the property of RFCs of assigning a continuous score in the [0.0,1.0] range to each data point. Thus, the decision threshold (which is at 0.5 by default) can be calibrated for samples from new sites using a small subset of manually rated data points. We can support this claim on the observation that the predicted “probability” of the RFC was close but below the default threshold of 0.5 (in the binary classification problem) for many of the misclassified data points of the held-out set *DS030*.

A second limitation of this work is the vague definition of MRI quality in our pre-registered report, which is closely related to the lack of agreement on how to grade the quality of images within the neuroimaging community. Instructing the experts with more detailed information on how to rate the images would have likely reduced the inter-rater variability and consequently the class-noise level. The labeling protocol presented here is very fast for the experts to visualize many images, but it is prone to class-noise as demonstrated by a fairly high inter-rater variability. In the early version of this manuscript, we used a quality assessment of ABIDE done by one of our experts with a different protocol. The change of protocol severely impacted the ratings (see SI: Impact of the labeling protocol and variability sources) and the performance evaluated on the held-out set due to the inconsistency of labeling protocols. An additional limitation of our labeling protocol is the use of reconstructed surfaces to aid raters. This approach introduces a bias in their judgment that would turn the general quality assessment into an evaluation of the particular tool used in the reconstruction (*FreeSurfer*). Therefore, the labeling protocol could be improved adding more resources to the rating settings (like the possibility of toggling the visualization of surfaces on and off, or the addition of visual reports generated from other processing tools or MRIQC itself). The raters do not pinpoint localized surface errors when no general defect is identified as their cause, in order not to bias their rating towards the evaluation of the reconstruction outcomes instead of the overall quality. Along the same lines, MRIQC does not include *FreeSurfer* in the extraction of IQMs to prevent leaking its performance into the features.

The spatial distribution of artifacts versus the global quality rating is another future line of research and current limitation. For example, the local motion of the eyes typically generates signal leakage through the phase-encoding axis. If the phase-encoding axis is anterior-posterior as opposed to left-right, the degradation is substantially more troublesome since the spillover will affect the lower regions of the occipital lobes (see Figure SI1). Future extensions of MRIQC should include the regional localization of the current IQMs. The feasibility of this approach is probably limited by the design principle of minimal preprocessing. Alternatively, the presented unsupervised framework could be replaced by a deep learning solution where the feature extraction is part of the design, and localization of quality features can be trained. We will also explore the integration of different modalities (e.g. T2-weighted). For instance, Alexander-Bloch et al. [4] propose the use of head motion estimated on same-subject, functional MRI time series as a proxy measure for motion during the T1w acquisition.

The quantitative assessment of quality using the RFC is the central piece of the three-fold contribution of this paper. The first outcome of this study is the MRIQC toolbox, a set of open-source tools which compute quality features. Second, MRIQC generates interactive visual reports that allow further interpretation of the decisions made by the classifier. Finally we propose the automated quality control tool described above to generate include/exclude decisions. We publicly release all the source code, the *Singularity* images and two classifiers to ensure the repeatability and transparency our experiments (see Software and data availability). Along with the tool, we release the quality ratings and all artifacts derived from training and testing the classifier to allow researchers to build upon our results or develop their own alternatives. For example, the quality ratings will allow MRI practitioners to train the model on a subset of their images and use a version of it *customized* for their site.

The MRIQC toolbox is a fork of the Quality Assessment Protocol (QAP). Since MRIQC was started as a standalone project, the implementation of most of the IQMs has been revised, and some are supported with unit tests. As with QAP, MRIQC also implements a functional MRI (fMRI) workflow to extract IQMs and generate their corresponding visual reports. Some new IQMs have been added (for instance, the CJV, those features measuring the INU artifacts, or the rPVEs). The group and individual reports for structural and functional data are also new contributions to MRIQC with respect to the fork from QAP. The last diverging feature of MRIQC with respect to QAP is the automated QC framework.

MRIQC is one effort to standardize methodologies that make data-driven and objective QC decisions. Automated QC can provide unbiased exclusion criteria for neuroimaging studies, helping avoid “cherry-picking” of data. A second potential application is the use of automated QC predictions as data descriptors to support the recently born “data papers” track of many journals and public databases like OpenfMRI [46]. For instance, MRIQC is currently available in the *OpenNeuro* [47] platform. The ultimate goal of the proposed classifier is its inclusion in automatic QC protocols, before image processing and analysis. Ideally, minimizing the run time of MRIQC, the extraction and classification process could be streamlined in the acquisition, allowing for the immediate repetition of ruled out scans. Integrating MRIQC in our research workflow allowed us to adjust reconstruction methodologies, tweak the instructions given to the participant during scanning, and minimize the time required to visually assess one image with the visual reports.

## Conclusion

The automatic quality control of MRI scans and the implementation of tools to assist the visual assessment of individual images are in high demand for neuroimaging research. This paper partially confirmed a pre-registered hypothesis about the feasibility of automated binary classification (“exclude”/”accept”) of the overall quality of MRI images. We trained a random forests classifier on a dataset acquired at 17 sites, and evaluated its performance on a held-out dataset from two unseen scanning centers. Classification performed similarly to previous works conducted on single-site samples. The hypothesis was not fully confirmed because we found that the classifier is still affected by a certain level of over-fitting to the sites used in training. Strategies aimed at combating site effects such as within site normalization and feature removal helped, but did not fully mitigate the problem. It is likely that adding labeled data from new sites will eventually ease this problem. We release all the tools open-source, along with the labels used in training and evaluation, the best performing classifier and all the derivatives of this work to allow researchers to improve its prediction and build alternative models upon this work.

## Author contributions

OE lead the development of MRIQC, implemented the cross-validation workflow, pre-registered the report, drafted the manuscript, run the experiments and interpreted the results. DB rated 601 data points of the ABIDE dataset and the *DS030* dataset in full. MS rated 601 data points of the ABIDE dataset, helped understand the problems of inter- and intra-rater variabilities. OOK contributed in the design of the cross-validation workflow, pre-registered the report and interpreted the results. RAP devised and coordinated the project, advised in all aspects of MRIQC, the cross-validation workflow and the manuscript design, pre-registered the report and interpreted the results. KJG devised the machine learning approach to quality control, coordinated the project, contributed to MRIQC and the cross-validation workflow, pre-registered the report, and interpreted the results. All the authors have read and edited the manuscript.

## Software and data availability

The pre-registered report is available online at https://osf.io/haf97/.

MRIQC is available under the BSD 3-clause license. Source code is publicly accessible through GitHub (https://github.com/poldracklab/mriqc). We provide four different installation options: 1) using the source code downloaded from the GitHub repository; 2) using the PyPI distribution system of Python; 3) using the poldracklab/mriqc Docker image; or 4) using BIDS-Apps [28]. For detailed information on installation and the user guide, please access http://mriqc.rtfd.io.

Two distributable classifiers are released. The first classifier was trained on ABIDE only, and it is the result of the experiment presented in Model evaluation and selection. The second classifier was trained on all the available data (including the full-ABIDE and the *DS030* datasets) for prediction on new datasets. Along with the tool, we release the quality ratings and all artifacts derived from training and testing the classifier. The animations used by the experts to rate the images are available at the pre-registration website. The *Singularity* images utilized in all the experiments presented here have been deposited to the Stanford Digital Repository (https://purl.stanford.edu/fr894kt7780).

The ABIDE dataset is available at http://fcon1000.projects.nitrc.org/indi/abide/. The *DS030* dataset is available at https://openfmri.org/dataset/ds000030/.

MRIQC can be run via a web interface without the need to install any software using OpenNeuro [47].

## Acknowledgments

This work was supported by the Laura and John Arnold Foundation. MS was supported by Swiss National Science Foundation (SNSF) grants (#158831 and 163859). The authors want to thank the QAP developers (C. Craddock, S. Giavasis, D. Clark, Z. Shehzad, and J. Pellman) for the initial base of code which MRIQC was forked from, W. Triplett and CA. Moodie for their initial contributions with bugfixes and documentation, and J. Varada for his contributions to the source code. CJ. Markiewicz contributed code and reviewed the second draft of the manuscript. JM. Shine and PG. Bissett reviewed the first draft of this manuscript, and helped debug early versions of MRIQC. S. Bhogawar, J. Durnez, I. Eisenberg and JB. Wexler routinely use and help debug the tool. We thank S. Ghosh for suggesting (and providing the code) the feature selection based on Winnow, and M. Goncalves for running MRIQC on their dataset, helping us characterize the *batch effect* problem. Finally, we also thank to the many MRIQC users that so far have contributed to improve the tool, and particularly to S. Frei who took the time to rerun our experiments.

